# Concurrent stimulation of diflufenican biodegradation and changes in the active microbiome in gravel revealed by Total RNA

**DOI:** 10.1101/2023.08.31.555673

**Authors:** Lea Ellegaard-Jensen, Pedro N. Carvalho, Muhammad Zohaib Anwar, Morten Dencker Schostag, Kai Bester, Carsten Suhr Jacobsen

## Abstract

The use of slowly degrading pesticides poses a particular problem when these are applied to urban areas such as gravel paths. The urban gravel provides an environment very different from agricultural soils; i.e., it is both lower in carbon and microbial activity. We, therefore, endeavoured to stimulate the degradation of the pesticide diflufenican added to an urban gravel microcosm amended with dry alfalfa to increase microbial activity.

In the present study, the formation of the primary diflufenican metabolite 2-[3-(Trifluoromethyl)phenoxy]nicotinic acid (commonly abbreviated as AE-B) was stimulated by the alfalfa amendment. The concurrent changes of the active microbial communities within the gravel were explored using shotgun metatranscriptomic sequencing of ribosomal RNA and messenger RNA. Our results showed, that while the active microbial communities in the gravel were dominated by bacteria with a relative abundance of 87.0 – 98.5 %, the eukaryotic groups, fungi and micro-eukaryotes, both had a 4-5 fold increase in relative abundance over time in the alfalfa amended treatment. Specifically, the relative abundance of microorganisms involved in degradation of complex carbon sources, Bacteroidetes, Verrucomicrobia, Sordariomycetes, Mortierellales, and Tremellales, were shown to increase in the alfalfa amended treatment. Further, the functional gene profile showed an increase in genes involved in increased activity and production of new biomass in the alfalfa treatment compared to the control, as well as pointing to genes potentially involved in biodegradation of complex carbon sources and the biotransformation of diflufenican.

## Introduction

Pesticides are applied not only on agricultural fields, but also in urban areas like public or private gardens, gravel paths, terraces, and courtyards. When pesticides are applied to urban gravel areas, their fate in terms of sorption, biodegradation, and leaching is often significantly different from their fate when applied on agricultural soils (Albers *et al*., 2020, Svendsen *et al*., 2020). Diflufenican is an example of an active ingredient in herbicide products applied in both agricultural and urban areas. Recent studies have found that diflufenican, which is used as a weedkiller in urbanised areas, was degraded much slower in urban gravel (e.g., used on gravel paths) than in agricultural soil, with no mineralisation detected and with the continuous formation of the metabolite AE-B (Svendsen *et al*., 2020). It was further shown that diflufenican sorbed much stronger in agricultural soil than in urban gravel, while the sorption of the metabolites AE-0 and AE-B was weaker than diflufenican in both soil and gravel (Svendsen *et al*., 2020). Together these findings suggest an increased risk of leaching of diflufenican and AE-B to groundwater when diflufenican is applied on gravel as compared to agricultural soils. Indeed, a leaching experiment with lysimeters columns, containing different gravel types and a sandy arable topsoil, showed that both of diflufenican’s degradation products (AE-0 and AE-B) leached to a greater degree from the gravel columns as compared to the soil columns (Albers *et al*., 2020).

The main difference between agricultural soil and urban gravel is the content of organic carbon (Svendsen *et al*., 2020), however key differences regarding the potential for diflufenican degradation are expected to be related to the microbial activity. As biotic degradation by microorganisms is chiefly responsible for pesticide transformation, previous studies have investigated the effects of pesticide application on soil microbial community and activity (Tejada, 2009, Baćmaga *et al*., 2015). A study applying a commercial herbicide containing diflufenican, mesosulfuron-methyl, and iodosulfuron-methyl-sodium in a pot experiment with sandy loam soil, found that the dose recommended by the manufacturer did not lead to significant changes in the size of the analyzed microbial populations determined as CFU’s of different microbial groups (Baćmaga *et al*., 2015). In the same study, however, an effect on the enzyme activity of the microbial community was reported (Baćmaga *et al*., 2015). Similarly, another study conducted over 180 days, found that diflufenican negatively affected the soil microbial biomass-C (fumigation-extraction) and enzymatic activities in two soil types (Tejada, 2009).

The slow degradation of diflufenican, with DT50 in lab studies (normalised) of 41.4-318 days and field studies DT50 224-621 days (Lewis *et al*., 2016, PPDB, accessed 3. July 2023), has led to studies attempting to stimulate its degradation by the addition of organic materials or biodegrading strains (Pinto *et al*., 2016, Carpio *et al*., 2020, Pinto *et al*., 2020). One such study added green compost to sandy loam soil without any significant increase in diflufenican degradation compared to the unamended soil (Carpio *et al*., 2020). Pinto *et al*. (2020) showed that sterile soil inoculated with three fungal strains *F. oxysporum, P. variotii*, and *T. viride* degraded 45 – 68 % diflufenican during a 120-day experiment. Thus, indicating that fungi may play a key role in degrading diflufenican *in situ*. These previous findings clearly highlight the need for further exploring the entire active microbial community in urban soils, e.g., gravel, and exploration of the potential role that soil fungi play in the transformation of diflufenican. Noticeably reports on the active microbiome in such systems are lacking in the literature.

In this context, the current study focused on investigating diflufenican biotransformation in an urban gravel, while examining the active microbial community using RNA Illumina shotgun sequencing and metatranscriptomic analysis of the entire active microbial community, i.e., of both prokaryotes and eukaryotes. Further, we applied a complex carbon source in the form of dried alfalfa to the gravel, which has been shown to increase bacterial and fungal soil biomass and activity (Mamilov *et al*., 2001). Hence this amendment was designed to stimulate the indigenous microbial community and thereby potentially enhance the biotransformation of diflufenican.

The objective of the present study was to stimulate microbial diflufenican transformation in an urban gravel soil matrix by adding a complex carbon source. We hypothesized that this would stimulate, especially, the microorganisms possessing the enzymes to break down complex organic compounds, thus concurrently stimulating diflufenican biotransformation.

## Materials and Methods

### Gravel

The type of gravel used in the microcosm experiment was of the brand *Slotsgrus* ™ (0 - 11.2 mm, Stenrand Grusgrav, Svebølle, Denmark). This gravel type is typically used in urban areas, where it is used for park trails and in courtyards in public and private areas. The gravel was sieved (<2 mm) and its physicochemical properties characterised previously by Svendsen *et al*. (2020).

Pellets of dry alfalfa (*Medicago sativa*)(Lucernepiller, dlg, Fredericia, Denmark) were triturated in a blender (JB3010WH, Braun, Braun GmbH, Germany) and mixed into one-half of the gravel at 0.2% (dw) immediately before the microcosm experiment (to simulate the microbial community by the release of carbon and nitrogen from the decaying plant litter), in accordance with previous application procedures (Mamilov *et al*., 2001, Schmitt *et al*., 2005, Johnsen *et al*., 2006).

### Chemicals

Diflufenican (N-(2,4-difluorophenyl)-2-[3-(trifluoromethyl)phenoxy]-3-pyridinecarbox-amide) was purchased from Sigma-Aldrich (Taufkirchen, Germany). Diflufenican metabolites AE B107137 (2-[3-(Trifluoromethyl)phenoxy]nicotinic acid; *abbreviation AE-B*) and AE 0542291 (2-[3-(trifluoromethyl)phenoxy]-nicotinamide; *abbreviation AE-0*) were acquired from Apollo Scientific (Cheshire, UK), and Chemieliva Pharmaceutical Co., Ltd, (Chongqing, China), respectively. Internal standard diflufenican-d3 was purchased from Toronto Research Chemicals (Toronto, Canada). Water (LiChrosolv LC-MS grade), methanol, and acetonitrile (LiChrosolv gradient grade) were acquired from Merck (Darmstadt, Germany). UltraPure Phenol:Chloroform:Isoamyl Alcohol in the ratio 25:24:1 used for RNA extaction was from Invitrogen (Thermo Fisher Scientific, Carlsbad, California).

### Microcosm experiment

Setup and sampling of the microcosm experiments were done in a laminar flow bench and all glassware was pre-sterilized by autoclavation. The microcosms were carried out in brown 60 mL glass jars. These were initially added 0.2 g sterilized Ottawa sand (50–70 mesh particle size, Sigma-Aldrich, Taufkirchen, Germany) each. Diflufenican dissolved in acetonitrile was added to the sand, where the acetonitrile was allowed to evaporate. Each jar was then added either 25 g (ww) alfalfa amended gravel or plain gravel and mixed thoroughly with the diflufenican-spiked sand to achieve a final concentration of 330 ng g^−1^.

This concentration is based on the recommended application dose 120 g ha^−1^ (Svendsen *et al*., 2020). Gravel water content was adjusted to pF = 2.2 immediately before use by the addition of sterile MilliQ water. The microcosms were covered with perforated aluminium foil and loosely closed lids to allow access to air. The experiment was incubated in darkness at 16 °C, and the water content was maintained by regular addition of sterile MilliQ water to compensate for any observed weight loss due to evaporation from the microcosms.

The microcosms were sampled once a month for chemical analysis over a six-month period (T0-T5). Here, 1.2 g samples were transferred from triplicate mesocosms of each treatment to 4 mL glass vials and frozen at -18 °C for subsequent processing. Likewise, samples for molecular microbial analysis (2 g) were transferred from the triplicate mesocosms of each treatment to sterile, RNase-free 15 ml Nunc tubes and immediately hereafter snap-frozen in liquid nitrogen and kept at -80 °C until RNA extraction. Samples from the timepoints T0, T1, T2, and T4 were selected for Total RNA library preparation (see section below).

### Chemical analysis of diflufenican and metabolites

Diflufenican and the metabolites AE-B and AE-0 were extracted from the gravel and analysed following Albers *et al*. (2020) and Svendsen *et al*. (2020). In brief; the samples were extracted by means of accelerated solvent extraction and quantified by HPLC-MS/MS. Limits of detection were 0.6 ng g^-1^ for diflufenican, 1.5 ng g^-1^ for AE-B, and 0.6 ng g^-1^ for AE-0, while recovery rates were 98, 76, and 75 %, respectively.

### Total RNA extraction, library prep and sequencing

RNA was extracted directly from the frozen samples with the addition of 2 ml of G2 DNA/RNA Enhancer (Ampliqon, Odense, Denmark) using the RNeasy PowerSoil Total RNA Kit (Qiagen, København, Denmark) with phenol:chloroform:isoamyl alcohol following the manufacturer’s instructions, except that the RNA was eluded in a final volume of 50 μl instead of 100 μl. DNase treatment was performed using DNase Max (Qiagen) according to the manufacturer’s instructions. The quantity and fragment size of the extracted RNA were determined on Qubit 4.0 fluorometer (Invitrogen, Eugene, Oregon, US) and Tapestation (Agilent, Santa Clara, CA, USA). The library was prepared using the NEBNext Ultra II Directional RNA Library Prep Kit for Illumina (New England BioLabs, Ipswich, MA, USA) in combination with the NEBNext Multiplex Oligos for Illumina, according to the manufacturer’s protocol. The fragment size of the resulting cDNA libraries was verified by Tapestation and DNA concentrations were measured on Qubit 4.0. Following, the samples were equimolarly pooled, and this final library was sequenced on an Illumina NextSeq 500 (Illumina Inc., San Diego, USA) with a High Output 300 cycles kit.

### Bioinformatics processing

In total, 9.8 × 10^8^ raw reads were obtained and processed through the quality control pipeline, as follows: Adapters were clipped using the TrimGalore (v0.6.6) tool; a wrapper script for automating cutadapt (Martin, 2011). FastP (Chen *et al*., 2018) was used for quality filtering of reads based on quality (q<20 in a window of 5 consecutive nts) and length (reads shorter than 60 nts) of the reads.

Prior to further analysis, reads were separated into small subunit (SSU) rRNA, large subunit (LSU) rRNA, and non-rRNA sequences through SortMeRNA (v4.3.1) (Kopylova *et al*., 2012). For the SSU, LSU, and non-rRNA read division across samples see Supplementary Table 1. The combined pool of SSU rRNA sequences from all samples was co-assembled into full-length SSU rRNA sequences using MetaRib (Xue *et al*., 2020). See Supplementary Table 1 for parameters other than the default ones. The assembly of rRNA reads was evaluated using QUAST (Gurevich *et al*., 2013). A total of 2.9 × 10^7^ ± 6.9 × 10^6^ reads for each sample were co-assembled into 3759 contigs. The N50 statistic of the assembly was 1472 with all 3759 contigs > 1000 nts. The contigs were taxonomically classified using CREST against the SILVA 128 database (Lanzén *et al*., 2012). SSU rRNA reads were mapped to the contigs using BWA (Li & Durbin, 2009) resulting in a table of taxonomically annotated read abundance across samples.

Stacked bar charts of the communities and NMDS plots based on Bray-Curtis dissimilarity were created in R (v4.1.1) (R-Core-Team, 2016) using the following packages and versions: phyloseq (v1.44.0) (McMurdie & Holmes, 2013), ggplot2 (v3.4.2) (Wickham, 2016), and tidyverse (v2.0.0) (Wickham *et al*., 2019). The vegan package (v2.6.4) (Oksanen *et al*., 2013) was used to run Adonis for multivariate analysis of differences between the communities of the treatments.

The combined pool of non-ribosomal sequences from all samples was processed through the CoMW pipeline (Anwar *et al*., 2019) to assemble complete mRNA sequences. Contigs were aligned against the M5nr protein database (Wilke *et al*., 2012), and annotated using eggNOG annotation. To identify the genes that were significantly differentially expressed between treatments, the DESeq2 (v1.34.0) (Love *et al*., 2014) module of the SARTools pipeline (v. 1.8.1) (Varet *et al*., 2016) was deployed in R using parametric mean–variance and independent filtering of false discoveries with the Benjamini–Hochberg procedure (p > 0.05) to adjust for type 1 error.

## Results

### Diflufenican degradation and metabolite formation

Samples from six different time points during the 150-day-long experiments were analysed for diflufenican and its metabolites. Diflufenican was only slowly degraded and no effect of alfalfa addition on the degradation of diflufenican (Figure 1A) was found. Accordingly, the final diflufenican concentrations in the alfalfa amended and control treatments of 189.2 (± 31.3) ng g^−1^ and 231.8 (± 25.1) ng g^−1^, respectively, did not differ significantly (final concentration t-test P= 0.140).

**Figure 1.**
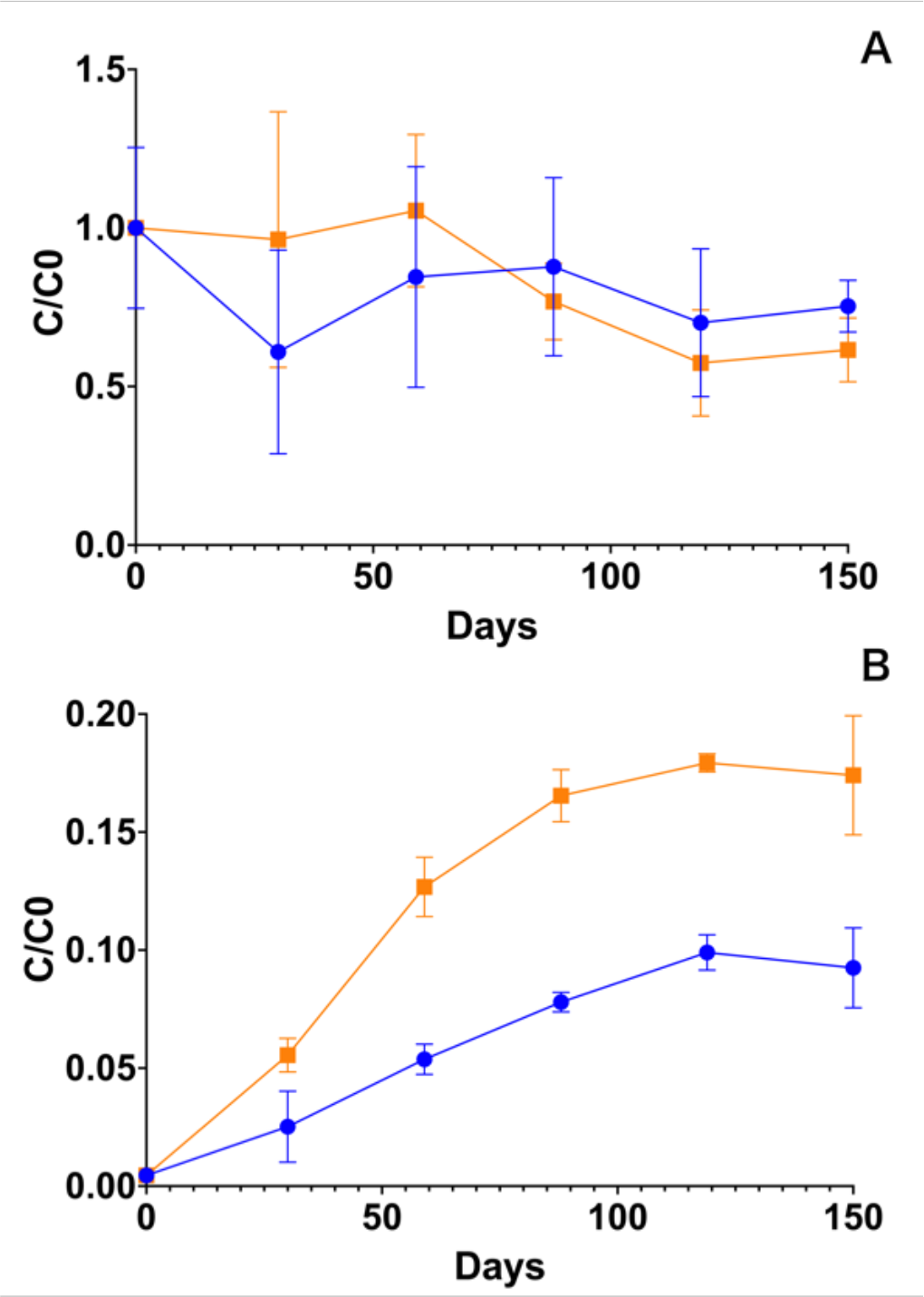
Pesticide residues in the treatments; alfalfa amendment (orange) and control (blue). (A) Degradation of diflufenican in the gravel as concentration/initial concentration (C/C0) as a function of time. (B) Formation of the metabolite AE-B from diflufenican spiked in gravel. The concentration/initial diflufenican concentration (C/C0) is calculated as the molar concentration of AE-B at time t, divided by the initial molar concentration of diflufenican at time t = 0. All points are averages of three replicate incubations with the standard deviation shown as error bar. AE-B stands for 2-[3-(Trifluoromethyl)phenoxy]nicotinic acid (EFSA ID - AE B107137).

However, in the alfalfa amended incubations higher concentrations of the metabolite AE-B were detected compared to the control as it was more rapidly produced and to a significantly higher final concentration (final concentration t-test P= 0.001; Figure 1B), with final concentrations of 38.5 (± 5.6) ng g^−1^ and 20.4 (± 3.7) ng g^−1^, respectively. In both treatments, AE-B concentration leveled out after 119 days. AE-0 was not detected in any of the samples.

### Total RNA sequencing

Samples from four different time points, day 0 (T0), 30 (T1), 59 (T2), and 119 (T4), were selected for Total RNA sequencing based on the chemical results. In total, 9.83 × 10^8^ raw reads were produced from these 24 samples. Resulting in each sample having 4.05 × 10^7^ ± 7.76 × 10^6^ reads (average ± standard deviation, *n* = 24) following quality filtration. Of these, sorting gave on average 69.8 % SSU, 20.4% LSU, and 9.8% non-rRNA (used in the mRNA pipeline). Details are given in Supplementary Table 1.

### Overall microbial community composition

A total of 3 734 distinct OTUs were found, of which 3 579 belonged to bacteria, 48 to SAR (Stramenopiles, Alveolates, and Rhizaria) supergroup, 44 to Amoebozoa, 39 to Opisthokonta, 14 to Archaea, 8 to Excavata, and 2 to Apusozoa (Figure 2B). Inspecting further the taxonomy of the eukaryotic groups and resorting these OTUs manually into ‘fungal’ and ‘other micro-eukaryotes’, 35 OTUs belong to fungal groups, while 106 belong to other Eukaryotic groups (i.e. micro-eukaryotes belonging to Amoebozoa, Apusozoa, Excavata, Opisthokonta, and SAR, but excluding fungal groups).

**Figure 2.**
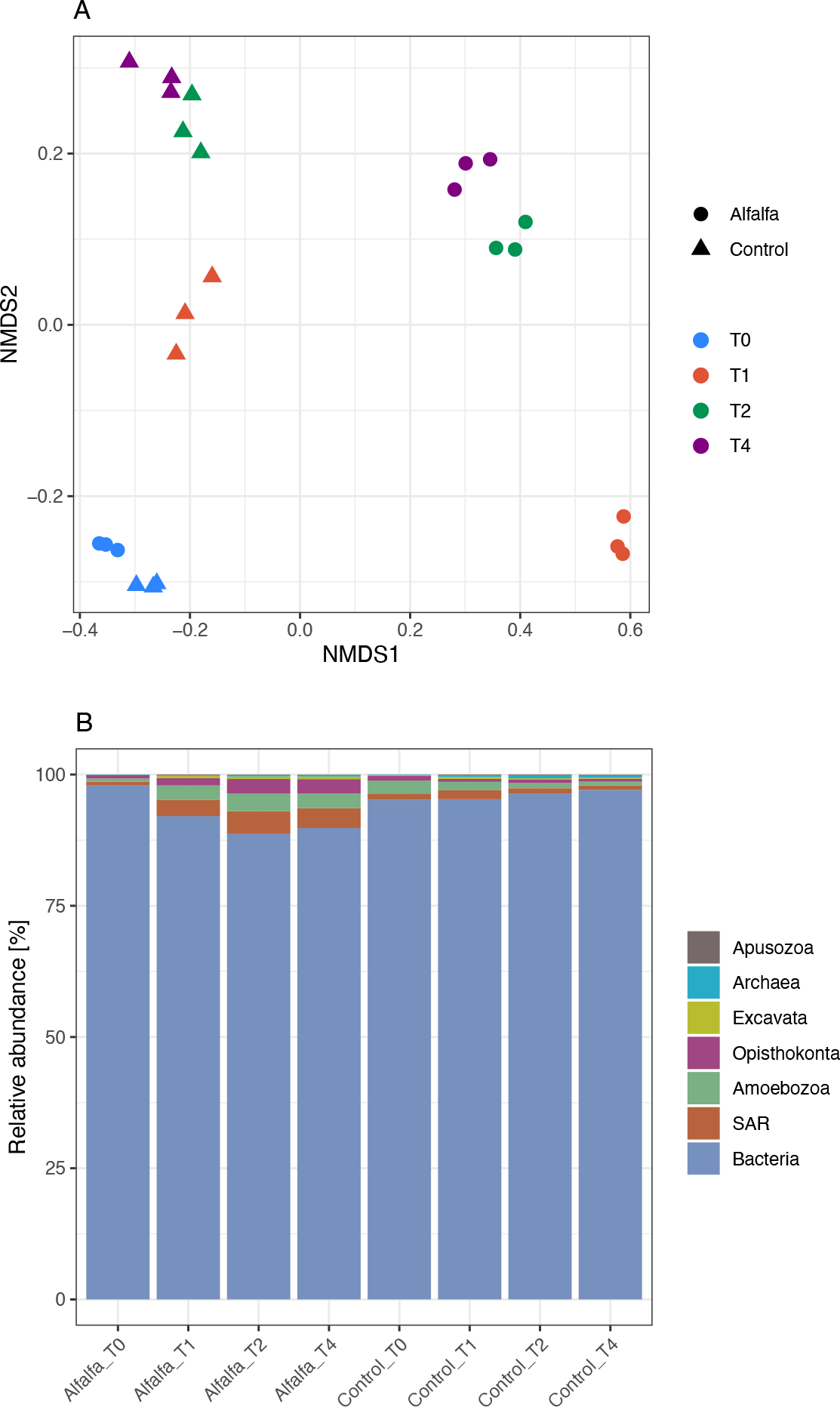
Changes in the overall community profile, based on the small subunit ribosomal RNA, for alfalfa amended and control treatments sampled at day 0 (T0), 30 (T1), 59 (T2), and 119 (T4). (A) Non-metric multidimensional scaling (NMDS) plot using Bray-Curtis dissimilarity matrices (stress = 0.0297, Shepard plot non-metric R 2 = 0.999) of the active microbial community (rRNA). (B) Composition of the entire active microbial community (rRNA) at Kingdom or supergroup level.

The community composition of the active microorganisms differed significantly between alfalfa amended and control treatments (p < 0.001, R^2^ = 0.28, *Adonis* (treatment) at OTU level, n = 24). The difference appears mainly driven by diverging communities at the three last time points (Figure 2A), and a significant difference is also found between the community composition of the different sample types i.e., of specific treatment and sampling time combinations (p < 0.001, R^2^ = 0.94, *Adonis* (treatment_timepoint)). Although the individual post hoc test was not significant for any two treatments (all p > 0.05, *Pairwise Adonis*).

Bacteria had by far the highest relative abundance in all samples constituting 87.0 – 98.5 % of the overall community (Figure 2B and Figure 3A). For alfalfa amended samples the relative abundance of bacteria decreased from T0 with 97.9 ± 0.2 % to T4 with 89.8 ± 0.8 %, while the relative abundance of bacteria in the control remained more stable within the range of 94.4 – 98.5 %. The relative abundance of archaea was very low ranging from 0.04 – 0.69 % of the overall community (Figure 3B). For alfalfa amended samples the relative abundance of archaea decreased from T0 (0.11 ± 0.02 %) to T1 (0.04 ± 0.00 %) followed by an increase for T2 and T4 (0.29 ± 0.05% and 0.32 ± 0.05 %, respectively). Whereas for the control the relative abundance of archaea increased steadily from T0 (0.17 ± 0.01 %) to T2 and T4 (0.57 ± 0.07 % and 0.47 ± 0.20 %) (Figure 3B).

**Figure 3.**
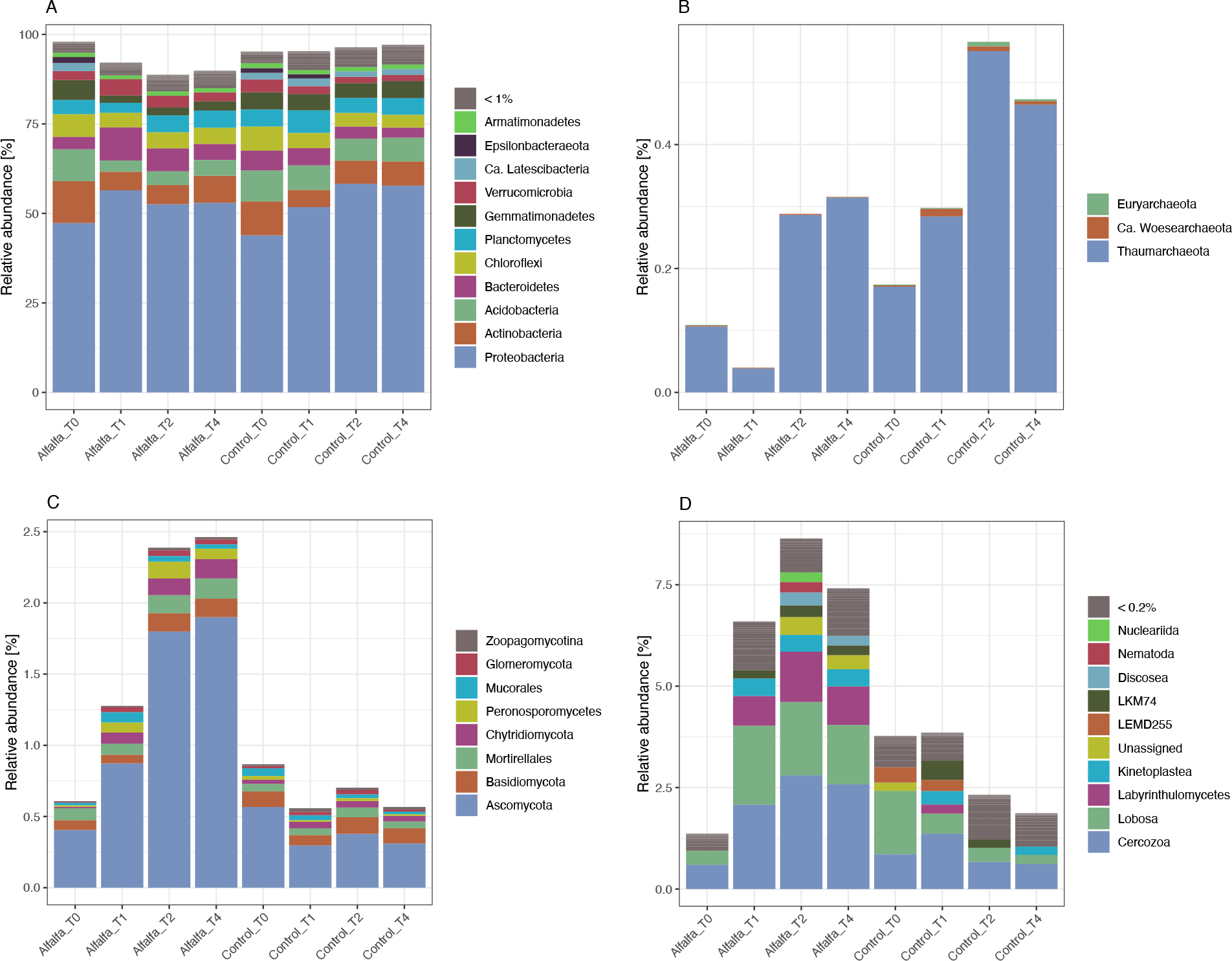
Composition of the active microbial community (rRNA) for alfalfa amended and control treatments sampled at day 0 (T0), 30 (T1), 59 (T2), and 119 (T4) divided into the following groups (A) Bacteria, (B) Archaea, (C) Fungi, and (D) Microeukaryotes.

The relative abundance of the combined fungal groups in the overall community ranged from 0.3 – 3.2 %. For alfalfa amended samples the relative abundance of fungi increased gradually from T0 (0.6 ± 0.1 %) to T4 (2.5 ± 0.2 %), whereas the abundance of fungi in the control samples remained relatively stable (0.7 ± 0.2 %) (Figure 3C).

The combined micro-eukaryotic taxa represented 0.9 – 9.6 % of the overall microbial community. An almost five times increase in relative abundance was seen in alfalfa amended samples from T0 with 1.4 ± 0.2 % to T1 with 6.6 ± 0.9 %, continuing to a maximum at T2 with 8.6 ± 0.9 % (Figure 3D). For the control samples, the relative abundance of the micro-eukaryotes was steady from T0 to T1 then it decreased slightly from T1 (3.9 ± 0.7 %) to T2 and T4 (2.3 ± 0.2 % and 1.9 ± 1.0 %) (Figure 3D).

### Prokaryotic community composition

The most abundant phyla of the bacterial communities were Proteobacteria > Actinobacteria > Acidobacteria > Bacteroidetes > Chloroflexi > Planctomycetes > Gemmatimonadetes > Verrucomicrobia (Figure 3A) as observed in order of descending relative abundance across all samples. Roughly the bacterial phyla can be divided into three groups; those that showed similar patterns for both treatments, those that showed an increase or decrease in relative abundance predominantly in the alfalfa amended samples, and those that show opposite patterns for the two treatments.

Proteobacteria as the most dominating among the bacterial phyla ranged from 42.8 – 62.4 % in relative abundance. For both the alfalfa amended and control samples the relative abundance of Proteobacteria increased from T0 (47.3 ± 2.0 % and 43.9 ± 0.9 %, respectively) to the following time points (T4; 53.0 ± 1.8 % and 57.8 ± 4.1 %, respectively). Actinobacteria also showed similar patterns for both treatments but with a decrease in relative abundance in both the alfalfa amended and control samples from app. 10 % at T0 to app. 5 %, 6 %, and 7% at T1, T2, and T4, respectively. While the relative abundance of Chloroflexi decreased in both treatments, but mostly in the control treatment where it decreased from 6.8 ± 0.3 % at T0 to 3.6 ± 0.2 % at T4.

The relative abundance of Gemmatimonadetes in the alfalfa amended samples decreased from app. 6 % at T0 to app. 2 % for the other time points, whereas it remained stable around 4 - 5 % in the control treatment. Acidobacteria decreased in relative abundance in both treatments, though to a larger degree in the alfalfa amended treatment from app. 9 % to between app. 3 and 4 % for T2 -T4. Bacteroidetes on the other hand showed an increase in relative abundance for the alfalfa amended samples only with 3.4 ± 0.4 % at T0 to 9.3 ± 0.7 % at T1 followed by a decline at T2 and T4. Whereas the control samples showed a steady decrease in the relative abundance of Bacteroidetes throughout all time points. Likewise, Verrucomicrobia showed an increase in relative abundance from T0 (2.5 ± 0.3 %) to T1 (4.6 ± 0.2 %) in the alfalfa amended samples, again followed by a decline at T2 and T4.

Lastly, the relative abundance of Planctomycetes decreased from T0 (3.9 ± 0.4 %) to T1 (2.7 ± 0.1 %) for the alfalfa amended treatment, while it at the same time increased for the control (4.7 ± 0.4 % to 6.3 ± 0.2 %). Hereafter, the relative abundance returned to app. 4 – 5 % for both treatments.

For archaea only three phyla were found. These were Thaumarcheotal > Ca. Woesearchaeota > Euryarchaeota (Figure 3B). Both Euryarchaeota and Ca. Woesearchaeota had relative abundances < 0.02 % at T0. They then increased to app. 0.1 % at some of the later timepoints for the control treatment, whereas they remained low for the alfalfa amended treatment. The relative abundance of Thaumarcheotal also showed the highest increase from T0 (0.17 ± 0.01 %) to T2 and T4 (0.55 ± 0.08 % and 0.46 ± 0.20 %) for the control treatment. While for alfalfa amended samples the relative abundance decreased from T0 (0.11 ± 0.02 %) to T1 (0.04 ± 0.00 %) followed by an increase for T2 and T4 (0.29 ± 0.05% and 0.31 ± 0.05 %, respectively).

### Eukaryotic community composition

The most abundant taxa of the fungal communities were Ascomycota > Basidiomycota > Mortirellales > Chytridiomycota > Peronosporomycetes > Mucorales (Figure 3C). All these taxonomic groups increased in relative abundance over time in the alfalfa amended treatment, while the relative abundances of these groups were either stable or decreasing in the control treatment over time. Ascomycota thus increased in relative abundance from T0 (0.41 ± 0.06 %) to T4 (1.90 ± 0.19 %) in the alfalfa amended treatment, while a slight drop from T0 (0.57 ± 0.07 %) to T4 (0.31 ± 0.13 %) was seen for this phylum in the control treatment. The rest of the taxa had a relative abundance < 0.2 % at all time points. Still, from T0 to T4 increases in relative abundance of between 1.5 – 13 times were observed in the alfalfa amended treatment for the rest of these phyla.

The remaining eukaryotes - i.e., the micro-eukaryotes within Amoebozoa, Apusozoa, Excavata, Opisthokonta, and SAR, excluding fungal groups – belong to nine taxonomically assigned groups with relative abundance above 0.2 % (Figure 3D). Across all samples they were observed in the following order of descending relative abundance Cercozoa > Lobosa > Labyrinthulomycetes > Kinetoplastea > LEMD255 > LKM74 > Discosea > Nematoda > Nucleariida. Eight of these groups being protists, while the last one was the phylum Nematoda (multicellular; Animalia). All but one of these groups increased in relative abundance from T0 to the following time points (LEMD255 being the exception) in the alfalfa amended treatment. While most of the groups either remained relatively stable or decreased in relative abundance in the control treatment. The Cercozoa, for instance, increased in relative abundance in both the alfalfa amended and control treatment from T0 (0.60 ± 0.05 % and 0.85 ± 0.08 %, respectively) to T1 (2.08 ± 0.19 % and 1.37 ± 0.20 %, respectively), followed by a further increase for the alfalfa amended treatment to 2.58 ± 0.25 % at T4, while the relative abundance decreased in the control treatment to 0.62 ± 0.33 % at T4.

### mRNA – functional genes

Annotation of the assembled contigs resulted in a combined 1009 COGs and NOGs across the samples. Comparing the alfalfa amended and control treatment at the four time points showed that the number of differentially expressed functional genes was highest at T1 with 230 genes differentially expressed between the two treatments. At the other three time points, there were approximately 100 differentially expressed genes between the two treatments.

A large proportion of the genes differentially expressed between the two treatments falls within the category ‘Poorly characterised’ e.g., with 108 within ‘Function unknown’ and 18 within ‘General function prediction only’ at T1 (Figure 4 and Supplementary Figure S1). Within the gene categories of the better-characterized genes especially ‘Translation, ribosomal structure and biogenesis’, ‘Transcription’, ‘Signal transduction mechanisms’, and ‘Posttranslational modification, protein turnover, chaperones’ showed elevation in the number of COGs/ NOGs which are more highly expressed in the alfalfa amended treatment than the control at timepoint T1 compared to the other time points. Likewise, the category ‘Carbohydrate transport and metabolism’, which could contain genes relevant for the breakdown of organic compounds (e.g., ‘COG0726 Predicted xylanase/chitin deacetylase’), showed an increase in the number of differentially expressed genes in the alfalfa amended treatment than the control at timepoint T1 compared to the other time points.

**Figure 4.**
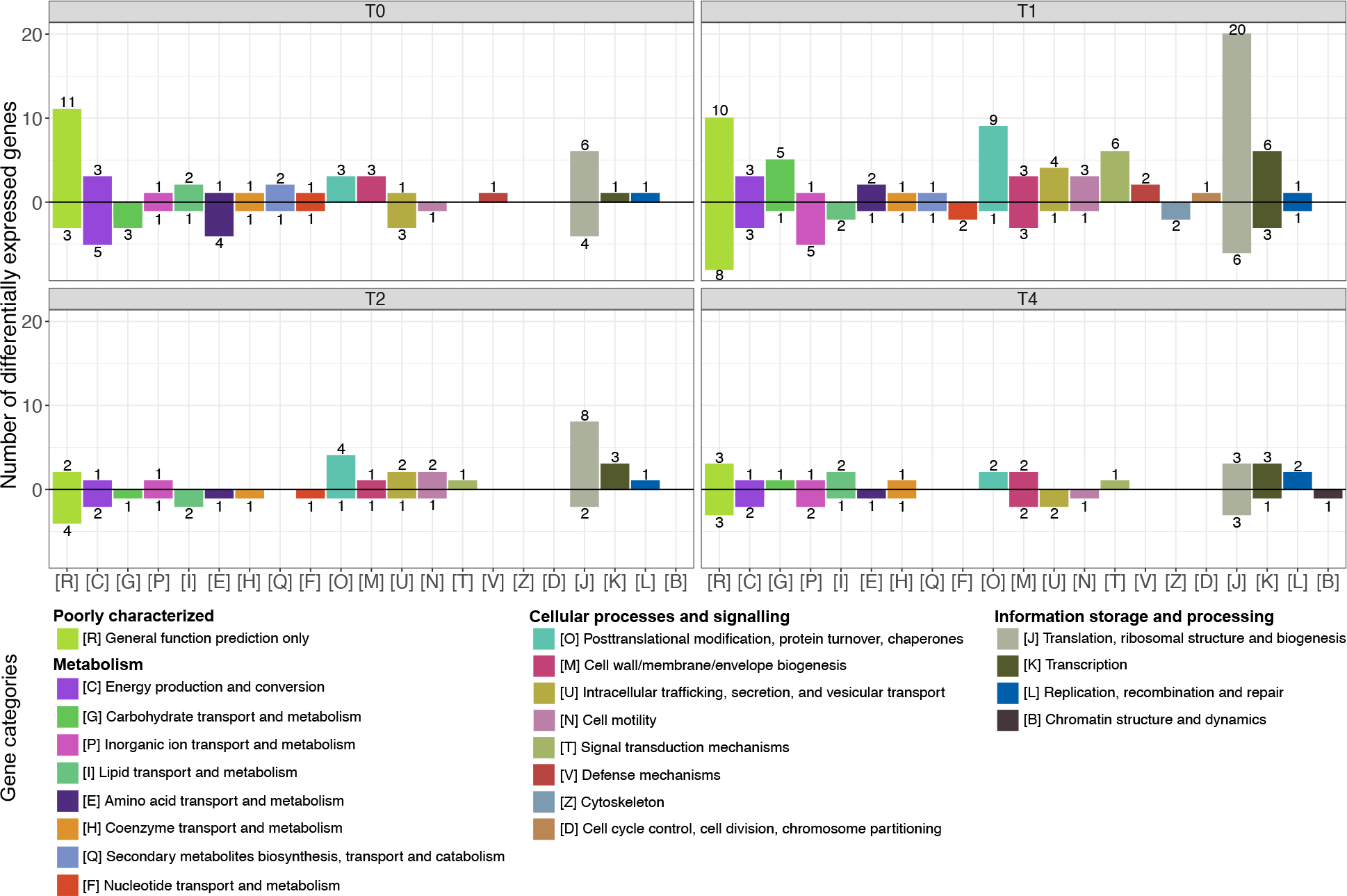
Numbers of differentially expressed genes within functional categories across alfalfa amended and control treatment by pairwise comparisons of gene transcription levels between samples at different incubation times; ‘T0’, ‘T1’, ‘T2’ and ‘T4’. Increasing and decreasing gene transcription levels are presented above and below the black horizontal zero line, respectively. Digits above/below bars represent the number of differentially expressed genes within a gene category. Gene category ‘[S] Function unknown’ is omitted. See Supplementary Figure S1 for all gene categories.

## Discussion

The biotransformation of semi-persistent pesticides applied to urban soils can be correlated to the organic carbon content of the soils (Meftaul *et al*., 2023). The reason for this being that the urban soils, e.g., gravel walkways, are often low in organic carbon and hence also low in microbial activity. In the current study, we therefore added organic carbon to urban gravel in order to stimulate the indigenous microbial community and thereby enhance the biotransformation of the pesticide diflufenican. The urban gravel used in the current study contained 0.17 % ***C***_***org***_ (Svendsen *et al*., 2020), which is comparable to other urban soils (Meftaul *et al*., 2023), and at least 10 times lower than that of an agricultural soil (Svendsen *et al*., 2020). The amendment of 0.2 % alfalfa (w/w) in the present study is comparable to what is used in other studies and roughly matches agricultural carbon application rates (Mamilov *et al*., 2001, Schmitt *et al*., 2005, Johnsen *et al*., 2006). In the present study we found that alfalfa amendment significantly stimulated the formation of the primary degradation product AE-B, with final concentrations corresponding to 17% and 9% of the added diflufenican for alfalfa and control treatment, respectively. However, this significant effect was only true for the metabolite formation, while the diflufenican removal was similar between treatments due to the large variation of diflufenican results.

Svendsen *et al*. (2020) previously reported 83% of the initial diflufenican remained in the same type of urban gravel after 150 days. In the present study 61% and 75% of the spiked diflufenican remained in the alfalfa amended and control treatments after 150 days, respectively. Although this suggests an enhanced diflufenican removal in the alfalfa amended gravel a clear conclusion cannot be made due to the large variations in the data. A larger standard deviation in diflufenican concentrations in gravel compared to soil due to was previously seen (Svendsen *et al*., 2020). Together this implies that soil degradation experiments protocols may need to optimised, e.g., sample mass or longer experimental duration, to accommodate for the higher variation resulting from urban gravel samples.

In comparison, we previously found only 43% of the initial diflufenican present after 150 days of incubation in a sandy agricultural soil (Svendsen *et al*., 2020). Another study found that on average 73% of the amount of diflufenican added to two agricultural soils (a sandy loam and a silt-loam) remained after 196 days (Bending *et al*., 2006). Hence, diflufenican is considered semi-persistent in most soil environments.

The formation of AE-B from diflufenican in the present study reached a maximum after 119 days, thus indicating that the metabolite might be further degraded. However, we did not detect AE-0 at any time, which is in agreement with our previous experiments in urban gravel (Svendsen *et al*., 2020). Concurrently, we previously found that AE-B could potentially be degraded in the same type of gravel to an uncharacterized metabolite (Svendsen *et al*., 2020), which may impact the formation pattern seen for AE-B. This suggests that further research is needed to determine additional metabolites.

Thus, the alfalfa amendment clearly enhanced the AE-B formation and consequently also the diflufenican biotransformation in the urban gravel used present study. This is in line with previous studies showing that addition of ground lucerne (alfalfa) straw to the fine material of railway ballast (coarse texture and low organic matter content) stimulated microbial activity and led to increased biotransformation of diuron into the metabolites DCPMU and DCPU (Cederlund *et al*., 2007). Another study found no increase in diflufenican degradation when amending the soil with either a spent mushroom substrate or green compost in a field experiment where a herbicide containing diflufenican was applied (Carpio *et al*., 2020). Although the same study did find that the organic amendments had a stabilising effect on soil microbial biomass and structure, determined by the phospholipid fatty acid profiles, as well as on microbial dehydrogenase activity following herbicide application (Carpio *et al*., 2020). Collectively this supports our initial hypothesis that the addition of a complex carbon source has a stimulating effect on the biotransformation of semi-persistent pesticides in soils or gravel with low carbon contents and low microbial activity, like the urban gravel used in the present study.

The enhanced biotransformation was accompanied by a change in the composition of the active microbial community in the alfalfa amended treatment compared to the control treatment. The TotalRNA approach applied in the present study has the strength of revealing changes in the composition of the active microbial community across all domains of life as demonstrated in previous studies investigating the active microbiome of permafrost soil (Schostag *et al*., 2019), wood ash amended agricultural and forest soil (Bang-Andreasen *et al*., 2020), and perennial cave ice (Mondini *et al*., 2022).

In the present study, bacteria clearly dominated the active microbial community in the urban gravel. We saw a decrease in the overall relative abundance of bacteria over time in the alfalfa treatment, where the relative abundance of bacteria in the control was quite stable. Despite of this decrease in the relative abundance of the overall bacterial community in the alfalfa treatment, we did see an increase in the relative abundance of Bacteroidetes and Verrucomicrobia both of which are known to include species capable of degrading complex carbohydrate-based biomass (Dash *et al*., 2020, Larsbrink & McKee, 2020). Whether these were involved in the transformation of diflufenican was not possible to elude from the present study and the literature on pesticide degraders within these phyla appears wanting. Further research is therefore needed to determine possible pesticide biodegrading species within these phyla.

The relative abundance of the combined fungi increased gradually over time in the alfalfa amended treatment, whereas the abundance of fungi in the control samples remained low and relatively stable over time. Most of the observed increase was due to an increase in Ascomycota, specifically taxa within class Sordariomycetes had a more than a teen fold increase in relative abundance. Members of this class are present in many ecosystems either as endophytes, pathogens or saprotrophs involved in decomposition (Taylor *et al*., 2015). Fungal species *F. oxysporum, P. variotii*, and *T. viride* previously shown to degrade diflufenican during a 120-day experiment (Pinto *et al*., 2020) likewise belong to class Sordariomycetes. This suggests that the alfalfa amendment may increase the abundance of Ascomycota fungi involved in diflufenican degradation, though these three species were not found in the present study.

In addition, other fungi may contribute to the degradation of diflufenican for instance species within the orders of Mortierellales and Tremellales, where we saw a slight increase in the relative abundance of the taxa within these orders in the alfalfa amended treatment compared to the control, but these could not be assigned beyond order level. *Mortierella* species have previously been shown to degrade diuron (Ellegaard-Jensen *et al*., 2013). Similarly, white-rot Tremellales species are capable of degrading complex carbon sources e.g., lignin and aromatic pollutants due to their production of polysaccharide- and lignin-degrading enzymes (Kirk *et al*., 1992). When fungi degrade organic material, they are generally using extracellular enzymes to facilitate the breakdown of complex compounds such as lignin and cellulose. These enzymes are non-specific and thus capable of degrading other complex compounds, e.g. pesticides, in soil (Magan *et al*., 2010).

Micro-eukaryotes are not directly involved in biodegradation. However, the play a vital part in the understanding of the microbial community changes as they graze on especially bacteria (Rønn *et al*., 2002). In the present study, we saw a vast increase in the overall abundance micro-eukaryotes over time in the alfalfa amended treatment compared to the control. This increase is mainly driven by the protozoan groups Cercozoa, Lobosa, and Labyrinthulomycetes (Figure 3). A great diversity was found in the current study for both Lobosa (Amoebozoa) and Cercozoa with 20 and 34 different taxonomical taxa, respectively. These most likely thrive on an increased bacterial growth in the alfalfa amended treatment, as they are both mainly considered to be bacterivores (Smirnov *et al*., 2011, Fiore-Donno *et al*., 2019). Though we did not measure the microbial biomass, Mamilov *et al*. (2001) previously found that an amendment with one percent alfalfa meal to a soddy-podzolic soil increased both the fungal and bacterial biomass in a mesocosm-experiment.

Furthermore, Chromadorea (Nematoda) also increase in relative abundance over time in the alfalfa amended treatment. This is in agreement with previous studies reporting an increase in number of Nematodes in alfalfa amended soils (Mamilov *et al*., 2001). The two dominating orders Araeolaimida and Rhabditida both feed on prokaryotes (Yeates *et al*., 1993), and are thus considered to contribute further to the overall grazing pressure on the bacteria and archaea within the alfalfa amended treatment. As such the micro-eukaryotes collectively exert a top-down control of particularly bacteria, and possible bacterial degraders may therefore be overlooked if they are heavily grazed.

The functional gene profile showed that especially genes involved in ‘Translation, ribosomal structure and biogenesis’ were more highly expressed in the alfalfa amended treatment than the control at timepoint T1 compared to the other time points. This points to an increase in the activity and production of new microbial biomass in the alfalfa treatment compared to the control, which is expected given the addition of extra carbon to a rather deprived matrix. Further, the application of such a complex carbon source was hypothesized to stimulates the expression of genes coding for potent catabolic enzymes. We found that the gene category ‘Carbohydrate transport and metabolism’, containing genes for breakdown of organic compounds, had five genes that had a significantly increased expression in the alfalfa amended treatment at timepoint T1. This included a gene for predicted xylanase/chitin deacetylase, which is well in line with breakdown of hemicelluloses from alfalfa given xylanase’s role in hemicelluloses degradation (Várnai *et al*., 2011).

Some genes coding for oxygenases were also found, but not showing a significantly increased expression. For instance, a gene for aromatic ring-cleaving dioxygenase was equally expressed in alfalfa and control treatments from T1-T4. Ring-cleaving dioxygenases are of particular relevance due to their efficacy in the degradation of environmental aromatic pollutants (Vaillancourt *et al*., 2006) - such as diflufenican. Finally, a gene annotated as Glyoxalase/Bleomycin resistance protein/dioxygenase (General function prediction only) had a great increase in the alfalfa amended treatment with a ∼30 FoldChange compared to the control at T0. We suggest, that further studies should focus on elucidation the role of the abovementioned genes and enzymes in diflufenican degradation.

We conclude that the amendment with a complex carbon source does enhance biotransformation of the recalcitrant pesticide diflufenican in urban gravel. Concurrently, the TotalRNA approach revealed increases in the relative abundance of active microbial groups potentially involved in the degradation complex carbon sources and diflufenican as well as grazers. Finally, the functional gene profile provided insides into the microbial gene expressions – where genes involved in increased activity and production of new biomass as well as potentially involved in biodegradation of complex carbon sources and biotransformation of diflufenican are seen as particularly relevant for future research into biodegradation of persistent pesticides.

## Acknowledgements

This project was funded by The Danish EPA (contract number 667-00225), by Aarhus University Research Foundation (AUFF-E-2017-7-21), and by the Novo Nordisk Foundation (Grant number: NNF19SA0059360). MZA was supported by the European Union’s Horizon 2020 - Research and Innovation Framework Programme under the Marie Skłodowska-Curie project MicroArctic (grant no. 675546), the Michael Smith Health Research BC trainee fellowship, and the Canadian Institutes of Health Research fellowship. MDS was supported by by the Danish National Research Foundation to the Center for Microbial Secondary Metabolites (DNRF137).

## Supplementary data

**Supplementary Table 1:** Sequence stats during bioinformatic processing.

|  |  | 1. Quality filtering and sorting into rRNA and mRNA |  |  |  |  |
| --- | --- | --- | --- | --- | --- | --- |
| Alfalfa treatment (%) | Time | 1.1 NextSeq output | 1.2 TrimGalore | 1.3 SortMeRNA |  |  |
|  |  | # Raw reads | # Sequences after QC | # rRNA SSU | # rRNA LSU | # Unaligned (mRNA) |
| 0 | T0 | 36,784,804 | 36,308,298 | 25,118,520 | 7,288,936 | 3,900,842 |
| 0 | T0 | 37,436,962 | 36,966,622 | 26,896,918 | 6,967,254 | 3,102,450 |
| 0 | T0 | 42,163,548 | 41,792,642 | 29,043,266 | 8,497,752 | 4,251,624 |
| 0.2 | T0 | 34,139,688 | 33,395,778 | 19,477,330 | 8,687,238 | 5,231,210 |
| 0.2 | T0 | 38,538,926 | 38,213,782 | 27,544,320 | 5,330,716 | 5,338,746 |
| 0.2 | T0 | 39,199,248 | 39,006,558 | 29,473,284 | 5,804,364 | 3,728,910 |
| 0 | T1 | 34,511,138 | 34,020,132 | 20,214,152 | 9,737,076 | 4,068,904 |
| 0 | T1 | 36,807,416 | 36,578,136 | 26,859,846 | 5,801,854 | 3,916,436 |
| 0 | T1 | 41,835,646 | 40,862,834 | 27,840,210 | 7,677,632 | 5,344,992 |
| 0.2 | T1 | 59,762,880 | 59,104,628 | 42,771,614 | 13,319,554 | 3,013,460 |
| 0.2 | T1 | 58,310,354 | 57,659,298 | 42,815,000 | 12,050,172 | 2,794,126 |
| 0.2 | T1 | 47,247,214 | 46,751,626 | 35,594,400 | 8,942,178 | 2,215,048 |
| 0 | T2 | 35,977,712 | 35,722,208 | 21,463,200 | 11,537,738 | 2,721,270 |
| 0 | T2 | 39,003,624 | 38,761,864 | 29,407,430 | 5,335,744 | 4,018,690 |
| 0 | T2 | 24,239,572 | 22,981,930 | 15,510,588 | 4,242,658 | 3,228,684 |
| 0.2 | T2 | 44,151,126 | 43,675,492 | 31,795,050 | 8,888,324 | 2,992,118 |
| 0.2 | T2 | 42,180,746 | 41,773,726 | 29,968,788 | 8,657,736 | 3,147,202 |
| 0.2 | T2 | 50,029,618 | 49,497,226 | 35,478,234 | 10,831,668 | 3,187,324 |
| 0 | T4 | 41,245,756 | 40,965,732 | 30,911,064 | 6,565,374 | 3,489,294 |
| 0 | T4 | 34,140,556 | 33,786,814 | 19,907,550 | 8,185,358 | 5,693,906 |
| 0 | T4 | 33,570,334 | 33,302,742 | 20,848,558 | 7,746,120 | 4,708,064 |
| 0.2 | T4 | 44,431,560 | 44,186,390 | 32,634,600 | 8,397,504 | 3,154,286 |
| 0.2 | T4 | 42,576,666 | 42,334,844 | 30,905,724 | 8,161,920 | 3,267,200 |
| 0.2 | T4 | 45,081,744 | 44,789,640 | 31,781,028 | 9,420,352 | 3,588,260 |
| Total sequences |  | 983,366,838 | 972,438,942 | 684,260,674 | 198,075,222 | 90,103,046 |

**Supplementary Table 1 (part 2/2): Sequence stats during bioinformatic processing.**
|  | <u>2. rRNA assembly</u> | <u>3. mRNA assembly</u> |
| --- | --- | --- |
|  | <u>2.1 MetaRib assembly<sup>1</sup></u> | <u>3.1 Trinity assembly</u> |
|  | <u>Assembly stats<sup>2</sup></u><br>(complete contig pool) | <u>Assembly stats<sup>3</sup></u><br>(complete contig pool) |
| # contigs | 3,759 | 1,042,590 |
| # contigs (>= 0 bp) | 3,759 | 1,042,590 |
| # contigs (>= 1000 bp) | 3,759 | 12,384 |
| # contigs (>= 5000 bp) | 0 | 89 |
| # contigs (>= 10000 bp) | 0 | 0 |
| Largest contig | 1,995 | 11,518 |
| Total length | 5,458,975 | 322,430,901 |
| Total length (>= 0 bp) | 5,458,975 | 322,430,901 |
| Total length (>= 1000 bp) | 5,458,975 | 18,912,466 |
| Total length (>= 5000 bp) | 0 | 573,642 |
| Total length (>= 10000 bp) | 0 | 56,861 |
| N50 | 1,472 | 285 |
| N75 | 1,400 | 238 |
| L50 | 1,798 | 363,144 |
| L75 | 2,741 | 674,004 |
| GC (%) | 55.78 | 56.83 |
| Mapped (%) | 72.49 | 89.28 |
| Avg. coverage depth | 11,887 | 30 |
| Coverage >= 1x (%) | 99.35 | 97.20 |

More information on the bioinformatic processing can be found in the Materials and Methods section.
<sup>1</sup>from MetaRib.cfg file; [EMIRGE], EM\_PARA : --phred33 -l 151 -i 300 -s 75 -a 64 -n 25, EMIRGE references: SILVA\_138.1\_SSURef\_NR99.
<sup>2</sup> Assembly stats produced using QUAST tool of 'all.dedup.filtered'
<sup>3</sup> Assembly stats produced using QUAST tool of 'Trinity'

**Figure S1.**
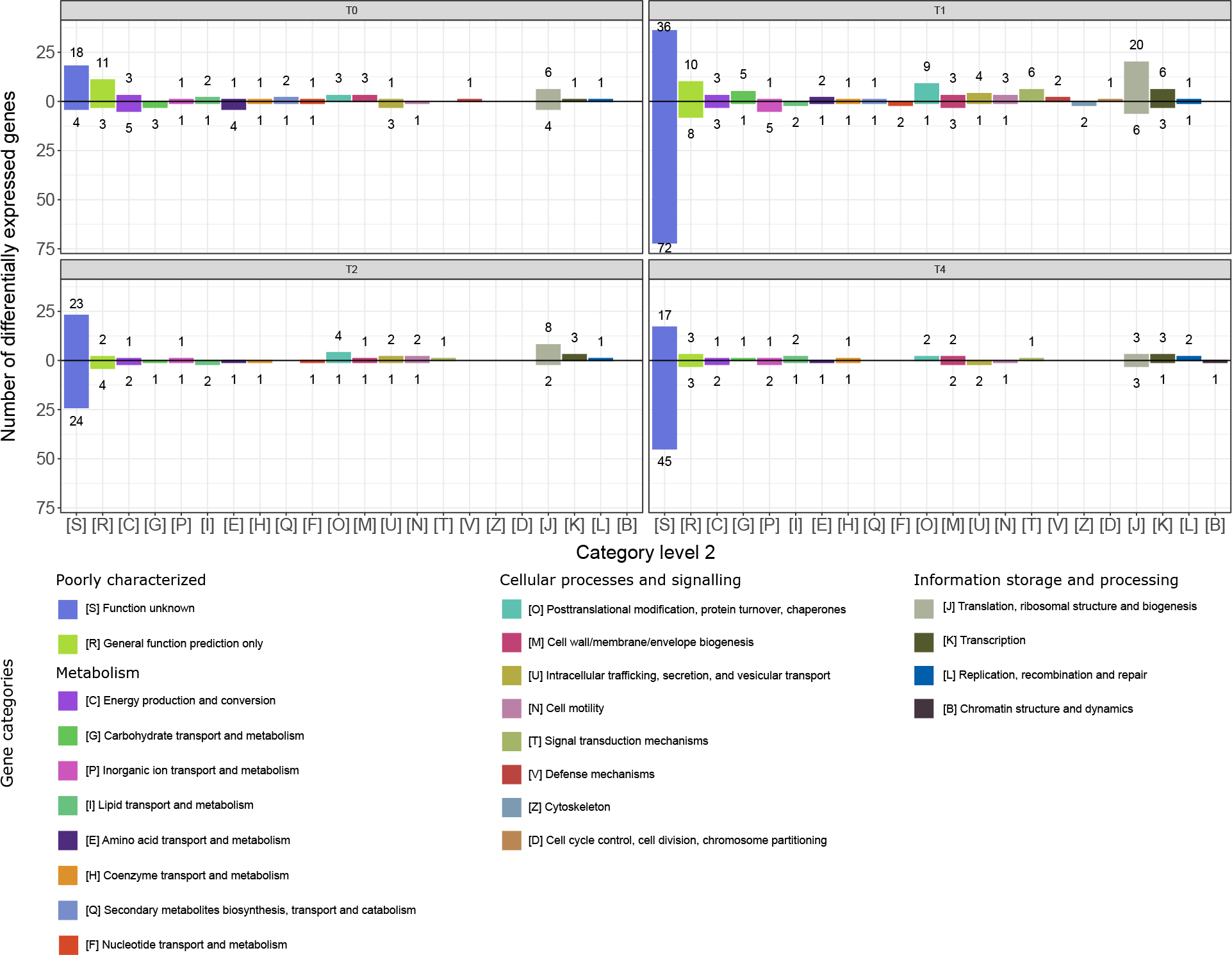
Numbers of differentially expressed genes within functional categories across alfalfa amended and control treatment by pairwise comparisons of gene transcription levels between samples at different incubation times; ‘T0’, ‘T1’, ‘T2’ and ‘T4’. Increasing and decreasing gene transcription levels are presented above and below the black horizontal zero line, respectively. Digits above/below bars represent the number of differentially expressed genes within a gene category.

